# A repressor-decay timer for robust temporal patterning in embryonic Drosophila neuroblast lineages

**DOI:** 10.1101/354969

**Authors:** Inna Averbukh, Sen-Lin Lai, Chris Q. Doe, Naama Barkai

**Author notes:** Co-first authors (Inna Averbukh, Sen-Lin Lai).

## Abstract

Biological timers synchronize patterning processes during embryonic development. In the *Drosophila* embryo, neural progenitors (neuroblasts; NBs) produce a sequence of unique neurons whose identities depend on the sequential expression of temporal transcription factors (TTFs). The stereotypy and precision of the NB lineages indicate reproducible temporal progression of the TTF timer. To examine the basis of this robustness, we combine theory and experiments. The TTF timer is commonly described as a relay of activators, but its regulatory circuit is also consistent with a repressor-decay timer, in which expression of each TTF begins once its repressor is sufficiently reduced. We find that repressor-decay timers are more robust to parameter variations compared to activator-relay timers. This suggests that the *in-vivo* TTF sequence progresses primarily by repressor-decay, a prediction that we support experimentally. Our results emphasize the role of robustness in the evolutionary design of patterning circuits.

## Introduction

Multicellular organisms shape their body plans during embryonic development through parallel processes that occur at different spatial positions and different times. Spatial coordination of patterning depends on direct cell-cell communication and long-range signaling by secreted morphogens. By contrast, temporal coordination is often achieved through cell-autonomous processes that measure time-delays (Pourquie, 1998). Molecular circuits implementing biological timers have been described (Murray, 2004; Reppert and Weaver, 2002; Simon et al., 2001), but the basis for their robust functioning is not well understood.

Biological timers play a key role in central nervous system (CNS) development of invertebrates and mammals, where a small pool of progenitors generates a vast amount of neuronal diversity (Grosskortenhaus et al., 2005; Isshiki et al., 2001; Kambadur et al., 1998; Kohwi and Doe, 2013; Kohwi et al., 2011; Li et al., 2013). In *Drosophila*, NBs delaminate from a ventral neuroectoderm at embryonic stages 9-11 to form an orthogonal two-dimensional grid with 30 NBs per half-segment. After formation, each NB undergoes asymmetric cell divisions every ˜45 minutes to produce a series of ganglion mother cells (GMCs), each of which divides into two post-mitotic neurons. The identity of each neuron is determined by the spatial position of the parental NB and by the TTF it inherits at the time of birth (Doe, 2017; Kohwi and Doe, 2013; Pearson and Doe, 2004). Temporal information is therefore conveyed by the NB cell-intrinsic timer, which drives the sequential expression of four TTF: Hunchback (Hb), Krüppel (Kr), Pdm (Flybase: Nubbin and Pdm2), and Castor (Cas) (Brody and Odenwald, 2000; Isshiki et al., 2001; Kambadur et al., 1998).

The molecular basis of the NB TTF timer has been previously characterized: Hb expression is initiated by an external signal, but subsequent dynamics depends on cross-regulation between the TTFs themselves (Cleary and Doe, 2006; Grosskortenhaus et al., 2006; Isshiki et al., 2001; Tran et al., 2010). In addition, the orphan nuclear hormone receptor, Seven-up (Svp), is required to switch off Hb expression, but not for specifying early neuronal fates (Kanai et al., 2005; Kohwi et al., 2011; Mettler et al., 2006; Tran et al., 2010).

The TTF expression sequence is largely independent of the cell cycle: Some NBs undergo just one cell division during the Hb expression window, whereas others undergo two or three divisions (Baumgardt et al., 2009; Doe, 2017; Isshiki et al., 2001). Furthermore, mutations that arrest the cell cycle still allow normal TTF expression from Kr onwards, and NBs can undergo the entire expression sequence when cultured in isolation in vitro (Brody and Odenwald, 2002; Grosskortenhaus et al., 2005; Kambadur et al., 1998). This indicates that the synchrony of the TTF timer with the cell cycle, needed to ensure reproducible NB lineage, is achieved by limiting the temporal variations in these two largely independent processes.

We hypothesize that reliable progression of the TTF cascade derives from the cell-intrinsic ability of the TTF timer to buffer variation (noise) in its molecular parameters. To identify the basis of this robustness, we used theoretical and experimental approaches. The TTF-timer is commonly described as a relay of activators (Doe, 2017; Rossi et al., 2017). Its regulatory circuit, however, contains also reactions compatible with a repressor-decay timer, in which TTFs progression depends on the decay of repressors. The relative contributions of the activator-relay and the repressor-decay interactions to TTF progression depend on unknown molecular parameters. To distinguish the *in-vivo* parameter values, we computationally screened millions of parameter sets for consistency with reported phenotypes and for the ability to buffer parameter variations. We find that decay timers are significantly more robust than relay timers. Based on that, we predicted and subsequently verified that *in-vivo* TTF expression timing depends primarily on repressor decay. We conclude that NB temporal patterning in *Drosophila* is driven by a robust timer primarily encoded by repressor decay, while activator relay plays a minor role.

## Results

### The TTF regulatory circuit combines activator-relay and repressor-decay Interactions

The sequential expression of TTFs within the dividing NB is commonly described as a relay of activators, in which each activator accumulates until reaching a threshold needed for inducing the next activator in the cascade (Doe, 2017; Rossi et al., 2017). Examining the TTF regulatory circuit, however, we noted that in addition to these activator-relay interactions, the regulatory circuit includes two additional interaction types: backward interactions, whereby a TTF inhibits the expression of a TTF upstream in the cascade, and repressor-decay interactions, whereby an upstream regulator represses the expression of downstream TTF **(Figure 1A-B)**. Therefore, at least in principle, the TTF timer can progress not only through a relay of activators, but also through decay of repressors, where target genes are induced once a repressor decays below some threshold level.

**Figure 1:**
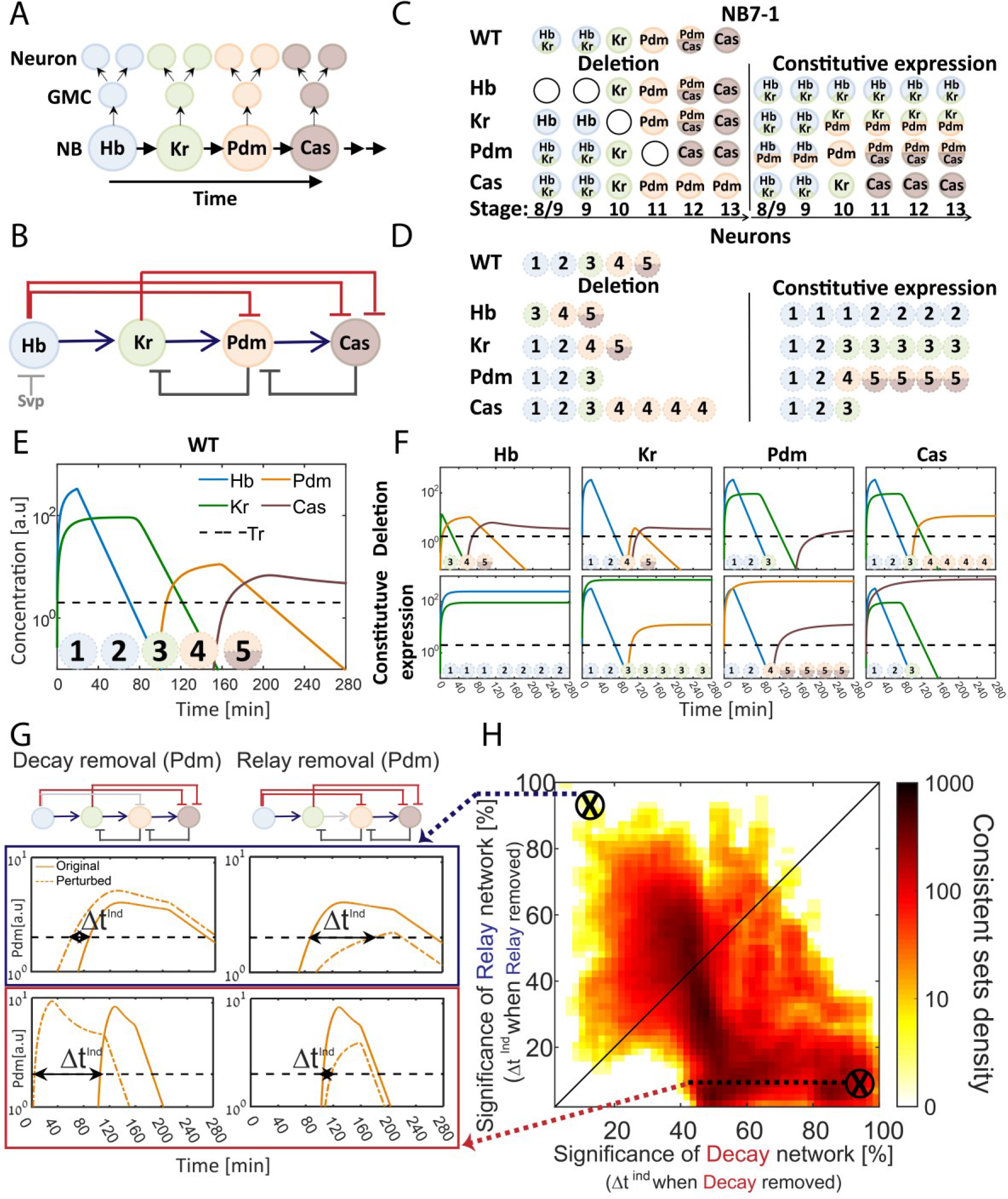
The TTF regulatory circuit combines activator-relay and repressor-decay timers. (A) The embryonic neuroblast TTF timer: In the *Drosophila* embryo, NBs express four TFs in a temporal sequence, as shown. We term this sequence the TTF timer. NBs divide asymmetrically to generate GMCs, whose further divisions produce post-mitotic neurons. The identity of both the GMCs and the post-mitotic neurons depends on the TTF expressed at the time of NB division, as shown in panel C, top left. (B) The TTF regulatory circuit: Experimentally defined cross-regulation between TTFs include interactions that propagate the cascade through activator relay (blue) or repressors decay (red), and backward interactions (black). (C) Summary of TTF expression in NB7-1 of wild type, mutant, and misexpression genotypes. TTFs are color-coded as in (A). Developmental stages indicated at the bottom and genotypes on the left. Data from (Grosskortenhaus et al., 2006; Isshiki et al., 2001; Tran et al., 2010). (D) Summary of neuronal identity in the NB7-1 lineage of wild type, mutant, and misexpression genotypes. Data from (Grosskortenhaus et al., 2006; Isshiki et al., 2001). (E) TTF timer model reproduces all reported phenotypes: A model was formulated that includes all regulatory interactions shown in Fig. 1B. Shown are the simulated dynamics of TTFs expression for a set of experimentally consistent parameters. Progeny identity was defined by the TTFs whose expression exceeded a constant threshold (dashed line) at the time of division (See **Table S1** for mapping of expressed TTFs to neuronal fate). (F) Mutant (top) and constitutive expression (bottom) models using the same parameter set as in 1E (see Methods for details). (G-H) Consistent circuits are distributed through the relay-decay timer space: over 10^6^ circuits differing by parameters choice were considered. A subset of ˜10^5^ circuits reproduced all experimentally defined phenotypes and are referred to as consistent circuits, or consistent parameter sets. Each consistent circuit was positioned on the relay-decay plot (H) based on the changes in Pdm and Cas induction (ind) times following removal of the respective activator-relay or repressor-decay interactions (grey bars, top of G). The density of consistent circuits in the relay-decay plot is color-coded. Note the larger density of consistent circuits in the regime of decay-timers. See Methods for details.

The activator-relay and repressor-decay reactions could both contribute to the progression of the TTF timer. Alternatively, one timer type could dominate. The relative contribution of the different reactions to the initiation of TTFs expression depends on the *in-vivo* parameters, whose values are not known and are difficult to measure. We therefore examined whether the *in-vivo* parameters can be distinguished computationally.

We formulated a model of the TTF timer that includes all experimentally described interactions, capturing their strengths by 17 independent parameters such as TTF production and degradation rates (Supplemental Information). The model is summarized by four ordinary differential equations (ODEs), whose solution simulates the temporal dynamics of the four TTFs. Expression thresholds define the minimal TTF expression levels required for transmission of this factor to the NB progenitors (Supplemental Information).

To define parameters consistent with the *in-vivo* dynamics, we compiled phenotypes of mutant embryos, focusing on the best characterized NB7-1 lineage **(Figure 1C-D).** Available data describes which TTF(s) are expressed by the dividing NB, within the GMCs, and by the postmitotic neurons. This data is of low temporal resolution, and cannot be used to define the precise durations at which each TTF is expressed. Still, this data provides the basic constraints with which our model should comply: the temporal sequence at which TTFs are expressed in our simulations should be consistent with the number of NB divisions, GMCs or post-mitotic neurons, reported for wild type, mutant, and misexpression embryos.

Screening over millions of parameters, each defining a different circuit, we detected a large number of parameter sets that were consistent with the reported phenotypes **(Figure 1E-F)**. To examine if these consistent sets favor an activator-relay or a repressor-decay timer, we focused on the last two TTFs (Pdm and Cas) for which activating and repressing interactions were described, and examined how their induction time changes when specifically removing either the activator-relay or the repressor-decay interaction (**Figure 1G**). These values - the changes in TTFs induction times when removing either the activator-relay or the repressor-decay interactions - positioned each consistent circuits within the relay-decay timer space (**Figure 1H)**, defining the significance of two respective timers in the circuit.

We observed a higher density of consistent circuits in the region of repressor-decay timers. However, a large number of consistent circuits were equally dependent on both the relay and the decay interactions, and some were driven primarily by relay **(Figure 1H)**. We conclude that the available data is not of sufficient resolution to distinguish whether the *in-vivo* parameters progress the TTF timer through an activator-relay or a repressor-decay timer.

### A repressor-decay timer is more robust than an activator-relay timer

To obtain better resolution, we next added robustness criteria to our model. In support of this, we had previously shown that time delays measured by protein decay are more robust than time delays measured by protein accumulation (Rappaport et al., 2005). The reason for this differential robustness is easily appreciated: the time at which a protein decays between two thresholds is only moderately (logarithmically) sensitive to the values of these thresholds **(Figure 2A-C)**. By contrast, the time to increase protein levels between two thresholds depends at least linearly, and typically significantly stronger, on thresholds values **(Figure 2A-C)**.

**Figure 2:**
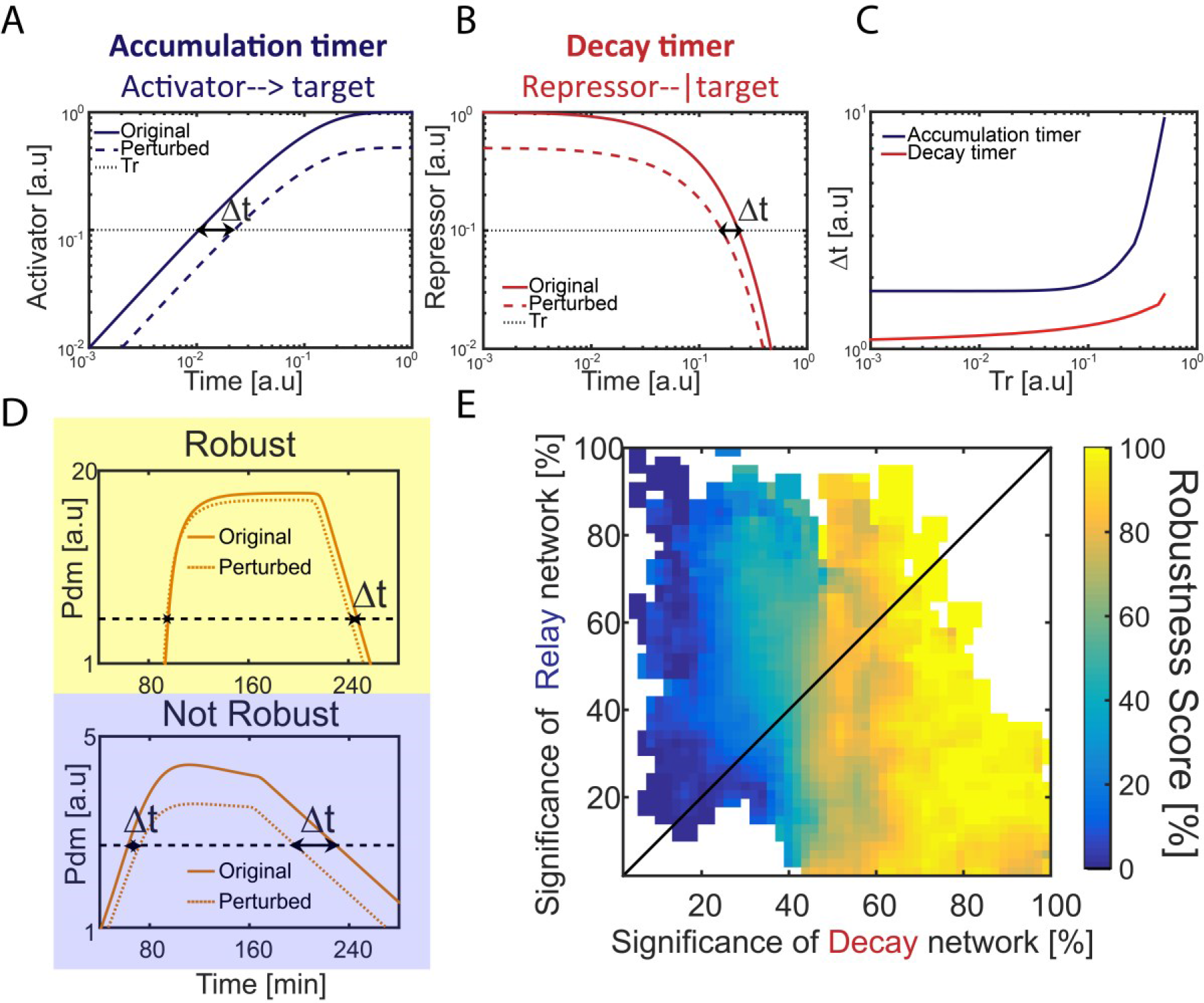
Repressor-decay timer is more robust than an 438 activator-relay timer. (A-C) Robustness of single-step timers: a single-step timer can be implemented by the accumulation of an activator (A) or by the decay of a repressor (B). In the accumulation of an activator scenario (A), activator production is initiated at t = 0. Once it accedes the threshold Tr, target genes are induced. In the decay of a repressor scenario (B), production of a repressor is stopped at t = 0. Once repressor levels have decayed below Tr, target genes would no longer be inhibited. The temporal dynamics of the regulatory proteins are shown in (A-B) for reference parameters (solid line) and following two-fold reduction in regulator production rate (dashed line). Timer output is defined by the time-delay from the onset of the dynamics until the regulator reaches the indicated threshold Tr. The change in this output following two-fold change in production rate is indicated (black double arrow), and is shown in (C) for different threshold values. See also analytical analysis in (Rappaport et al., 2005). (D-E) Distribution of robust circuits in the relay-decay timer space: consistent circuits, described in Fig. 1, were scored for robustness by measuring timer sensitivity to moderate (20%) variations in production parameters (see Methods). Examples for circuits showing different robustness scores are shown (D). Circuits were positioned on the relay-decay plot, as is Fig. 1H, and their robustness scores, averaged over closely-positioned circuits, are color-coded (E).

We added the robustness criteria to our numerical screen, assigning each consisted circuit a robustness score based on its ability to buffer the temporal durations at which each TTF is expressed against moderate (˜20%) variations in TTF production rates (see Methods) Examining the parameter sets that showed high robustness, we noted that they typically gave larger weights to the repressor-decay interactions compared to the activator-relay ones, suggesting that the more robust circuits progress the TTF timer through the repressor decay, rather than activator accumulation (data not shown).

To more rigorously distinguish whether robustness correlates with a specific timer type, we considered again the positioning of all circuits in the decay-relay timer space **(c.f. Figure 1H)**, and color-coded the circuits by their robustness score **(Figure 2D,E)**. High robustness scores were found in the region of repressor-decay timers, while activator-relay timers were significantly less robust. We conclude that also in the context of the full model, robustness is improved when progressing through repressor decay rather than activator relay.

### A TTF circuit can be positioned in the relay-decay timer space based on TTF-deletion phenotypes

We hypothesized that robust circuits, with improved ability to buffer variations in parameters, were favored in evolution. We therefore predicted that the *in-vivo* TTF timer is robust, and is thereby driven by a decay timer. To test this hypothesis, we searched for experiments that could distinguish properties of the *in-vivo* timer.

Experimentally, TTF deletion is the most accessible perturbation. As described above, the consequences of such perturbations were previously reported, but at a resolution that was too low to distinguish between the two timer types. Our simulations pointed to one limitation of existing data: for some mutants, the consequence of TTF deletion was defined by measuring the fates of the post-mitotic neurons, and therefore did not provide conclusive data about possible co-expression phases in which two consecutive TTFs are expressed within the NB, but one of them dominates in generating neuronal identity. This significantly limited our ability to precisely deduce the TTF expression timing in either wild-type or mutant embryos, and thereby greatly increased the spectrum of circuits that were scored as consistent with measured phenotypes.

With this in mind, we examined computationally whether the consequences of TTFs deletions, if analyzed at higher resolution, could distinguish between the repressor-decay and activator-relay timers. First, we examined how Pdm induction time changes following deletion of either Hb (its repressor) or Kr (its activator). Specifically, for each consistent circuit, as described in **Figure 1H** and **2E** above, we tested how Pdm induction time changes when simulating the removal of Hb and when simulating the removal of Kr **(Figure 3A)**. These two values allowed us to uniquely position each consistent circuit within the Hb-Kr sensitivity space **(Figure 3B).** Further, using the robustness value of each consistent circuit, we could color the consistent sets positioned on the Hb-Kr sensitivity space by their robustness score **(Figure 3B).** As can be appreciated, high robustness was found exclusively in circuits for which Pdm induction was dependent only on Hb decay. Analogous analysis of Cas induction time shows a similar, although less pronounced bias **(Figure S1)**. We conclude that measuring the change in Pdm induction time following Hb and Kr deletion can distinguish the decay Vs. relay properties of the *in-vivo* timer.

**Figure 3:**
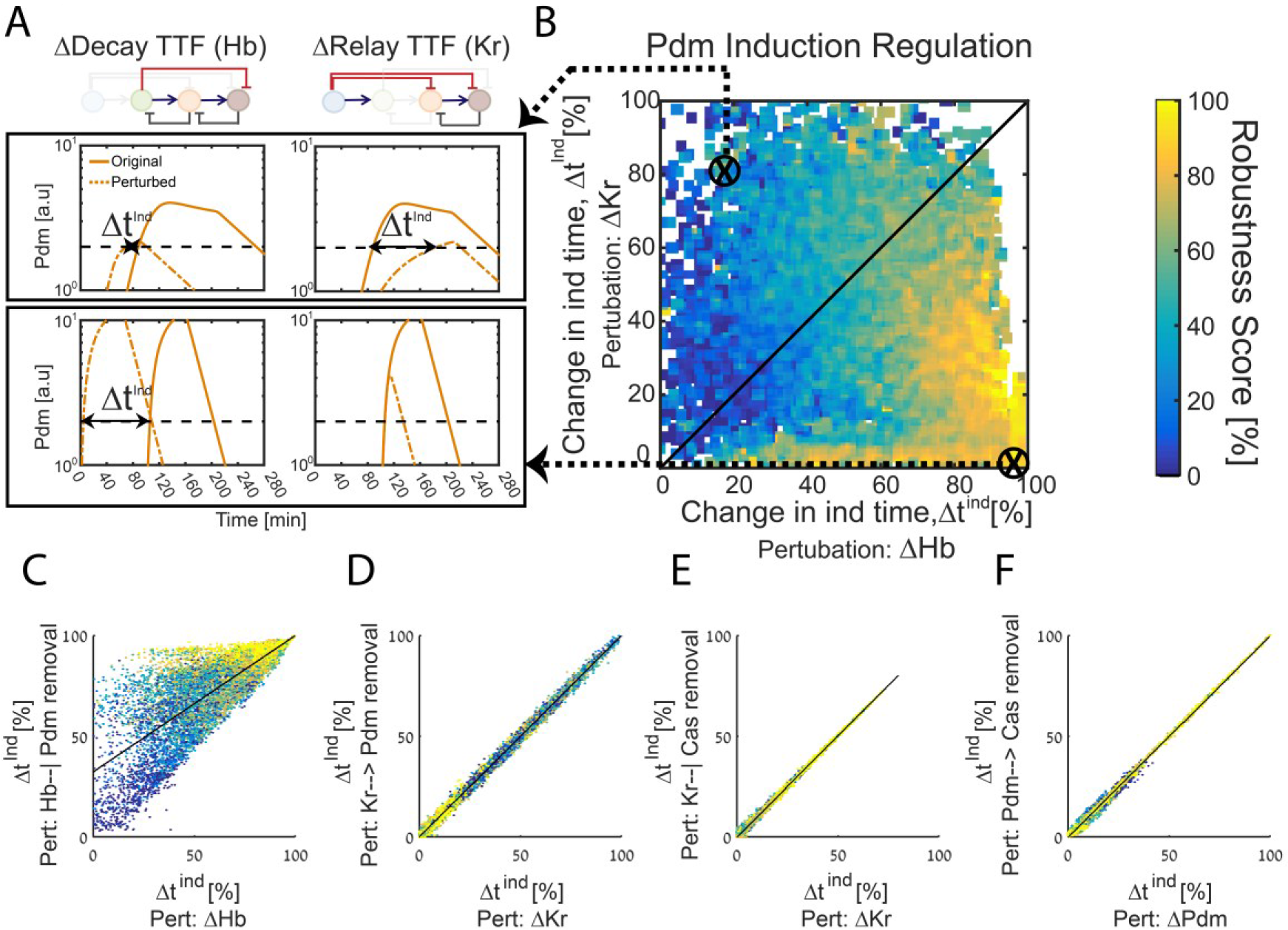
The TTF circuit can be positioned in the relay-decay timer space based on TTF deletion phenotypes. (A-B) TTFs deletion phenotypes can distinguish robust circuits: All consistent circuits, as described in Figure 1–2 above, were considered. Each consistent circuit was scored by measuring the change in Pdm induction times following deletion of Hb (A, left) or deletion of Kr (A, right). These values were used to uniquely position each circuit in the Kr-Hb sensitivity space (B). Color-coding circuits based on their robustness score, as in Figure 2E, shows that robust circuits are only found in a small region in the Kr-Hb sensitivity space, in which Pdm induction time is insensitive to Kr deletion. (C-F) Sensitivity to TTF deletion (X axis) correlates with the sensitivity to the specific removal of the respective activator-relay or inhibition delay interactions (Y axis): All consistent circuits, as described in Figure 1–2 above, were considered. For each consistent circuit, the changes in Pdm or Cas induction times following TTF deletion or removal of regulatory interactions was measured. Correlations between the effects of TTF deletion and removal of the respective regulatory link are shown. Each dot in these correlation figures represent one consistent circuit, color-coded by its robustness score.

Deletion of the TTF which functions as an activator or repressor abolishes the respective relay or delay interactions. However, it may have additional effects that are not directly related to these interactions. We therefore used our simulations to examine whether TTFs deletion phenotypes can predict the consequence of specifically abolishing the respective activator-relay or repressor-decay. To this end we considered again all consistent circuits. For each consistent circuit, we examined how Pdm induction time changes when specifically removing either the activator-relay (Kr-to-Pdm) or the inhibitor-decay (Hb-to-Pdm) interactions. This allowed us to compare, for each consistent circuit, the change in Pdm induction time when an upstream TTF (e.g. Kr) was deleted, or when the respective interaction (e.g. Kr-to-Pdm) was specifically removed **(Figure 3C,D)**. As can be seen, the consequences of these two perturbations were tightly correlated. A similar tight correlation was also observed when comparing the change in Cas induction time following the deletion of its activator (Pdm) or the removal of the Pdm-to-Cas activating link only, and when comparing the consequences of deleting the Cas repressor Kr to the specific removal of the Kr-to-Cas repression link (**Figure 3E,F)**. We conclude that following Pdm and Cas expression timing in the mutant embryos has the power to inform us not only about the robustness of the *in-vivo* timer, but also about the relative contributions of the relay and decay reactions to the TTF progression.

### Timing of Pdm and Cas expression is highly sensitive to deletion of TTF repressors, but less sensitive to deletion of TTF activators

As described above, our modeling suggests that activator-relay and repressor-decay timers can be distinguished based on the TTF deletion phenotypes, but that this would require data on TTF expression levels and timing at a much higher temporal resolution than had been obtained previously. To this end, we stained embryos for the TTF of interest, and for the Worniu and Engrailed markers, which allow us to unambiguously identify NB7-1 **(Figure 4A)**. TTF protein intensity levels were quantified using confocal microscopy **(Figure 4B)**. Variability in staining intensities was controlled by normalizing TTF staining to that of Engrailed, which is constantly expressed in NB7-1.

**Figure 4:**
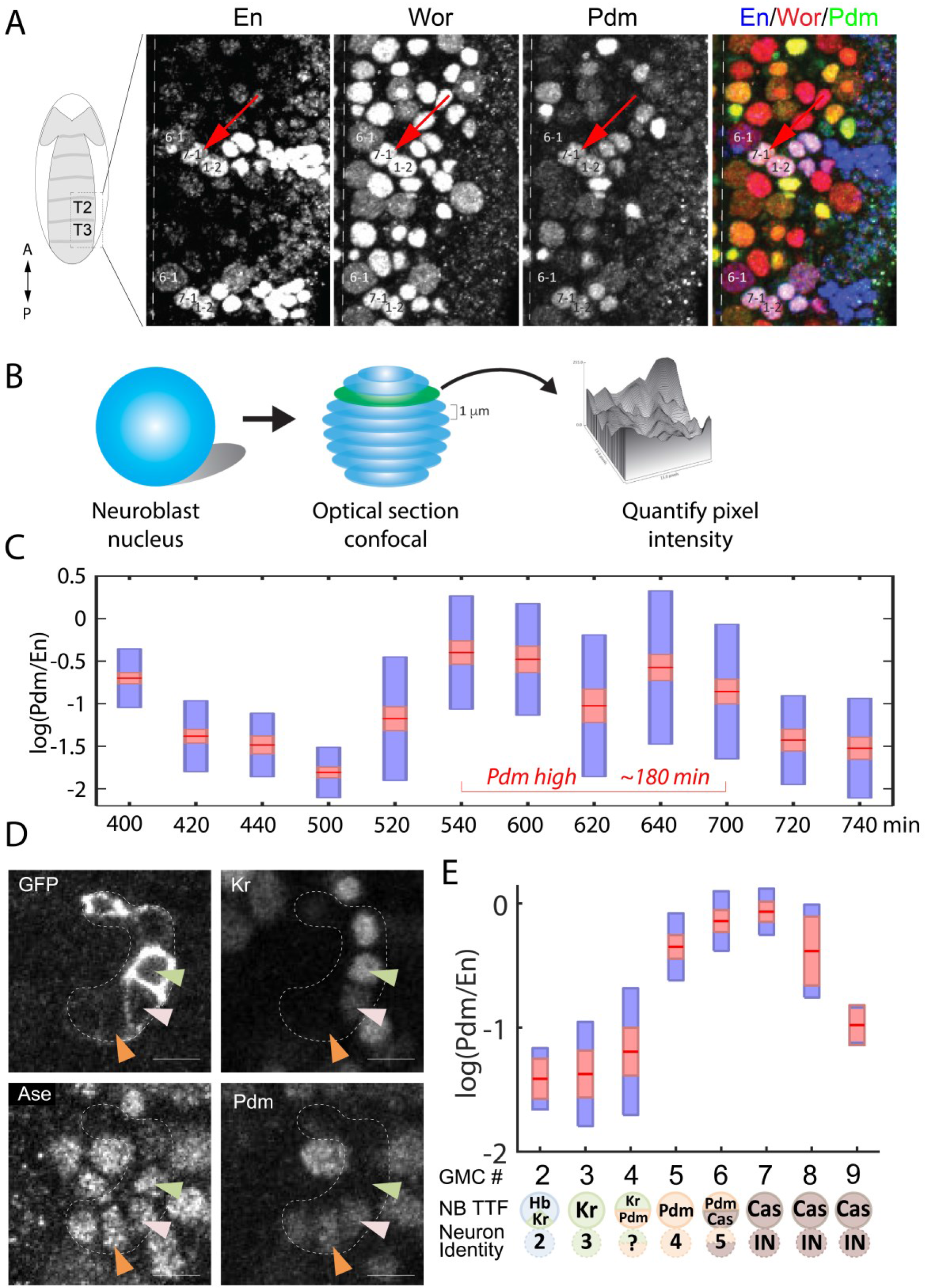
High temporal resolution analysis 480 of Pdm expression. (A) Confocal image of the neuroblast layer from ventral nerve cord segments T2 and T3 of an early stage 11 embryo (boxed area in illustrated embryo on the left). The NB7-1 was identified by the pan-NB marker Worniu (Wor) and the NB spatial marker Engrailed (En). NB6-1 is the most anterior medial En+ NB, with the En+ NB7-1 just posterior and lateral (red arrow), and En+ NB1-2 completing the diagonal. Genotype: *y*^1^ *w*^1^. Scale bar: 20 um. (B) Methodology for obtaining transcription factor levels in G1/G2/S-phase of NB7-1. Confocal section stacks (at a 1 um interval) of individual NB nuclei were obtained, and the area and signal intensity of each section were measured and summed to obtain the total intensity of TTFs. (C) Data from the number of NB7-1 indicated in Table S2 is summarized by box plots of measured log(Pdm/En) staining intensity as a function of time. The 1.96 SEM (95% confidence interval) is shown as a red rectangle with a horizontal red line for the mean, with a blue rectangle marking the limits of one standard deviation above and below the mean. Time duration of the Pdm phase (approximately 180 minutes) is indicated by a red double arrow. (D) Confocal image of a NB7-1 lineage marked with GFP (*NB7-1-Gal4*, *UAS-GFP*) in an early stage 11 embryo. Arrowheads indicate three consecutive GMCs (Ase^+^) which are Kr^+^Pdm^−^ (green), Kr^+^Pdm^+^ (pink), and Kr^−^Pdm^+^ (orange). Genotype: *ac-VP16*^*AD*^ *gsb-Gal4*^*DBD*^ *UAS-superfoldGFP*. Scale bar: 5 um. (E) Data from the number of NB7-1 indicated in Table S2 is summarized by box plots of measured log(Pdm/En) staining intensity as a function of the number of progeny GMC. The 1.96 SEM (95% confidence interval) is shows in red rectangle with a horizontal red line for the mean, one standard deviation above and below the mean in blue rectangle. Scheme below the X axis shows which neuronal progeny is hypothesized to be derived from the newest GMC.

Our data confirmed the sequential expression of Hb, Kr, Pdm, and Cas within the NB7-1 lineage (data not shown). It further revealed that Pdm expression was longer than expected: about 180 minutes **(Figure 4C)**. This is long enough to generate more than the two previously reported Pdm+ GMCs (Isshiki et al., 2001). To determine if there were additional Pdm+ GMCs in the lineage, we used the NB7-1-specific Gal4 driver to drive the expression of membrane-tethered superfold GFP and co-stained for Pdm and the GMC marker Asense **(Figure 4D).** We found that Pdm was upregulated when NB7-1 was producing the 4th GMC and downregulated after the NB generated the 7th GMC **(Figure 4E)**, indicating that four GMCs are produced within the Pdm expression window. Consistent with this finding, we observed a novel Kr+Pdm+ GMC in the lineage, which was not previously reported (Isshiki et al., 2001). The newly-discovered GMC may produce an Eve-negative motor neuron, interneuron, or undergo programmed cell death (see Discussion).

The Pdm expression window is therefore significantly longer than the duration inferred from the previous data used to calibrate our model. This difference in the timing of Pdm expression in wild-type embryos has no substantial effect on our model. Indeed, apart from minor quantitative differences, our main qualitative results, including the ability to distinguish consistent circuits and the differences between the robustness of decay and relay times, remained the same.

We next used the high temporal resolution expression data to determine whether the relay-timer or decay-timer could best account for Pdm and Cas expression timing in TTF mutant backgrounds. We found that *Kr* mutants did not alter Pdm expression, whereas *hb* mutants advanced Pdm by about two cell-cycles **(Figure 5A-C)**, showing that Pdm expression is more sensitive to deletion of the upstream repressor Hb. Similarly, *pdm* mutants did not have as much effect on Cas expression as did *Kr* mutants, showing that Cas expression is more sensitive to deletion of the upstream repressor Kr rather than the upstream activator Pdm **(Figure 5D-F)**.

**Figure 5:**
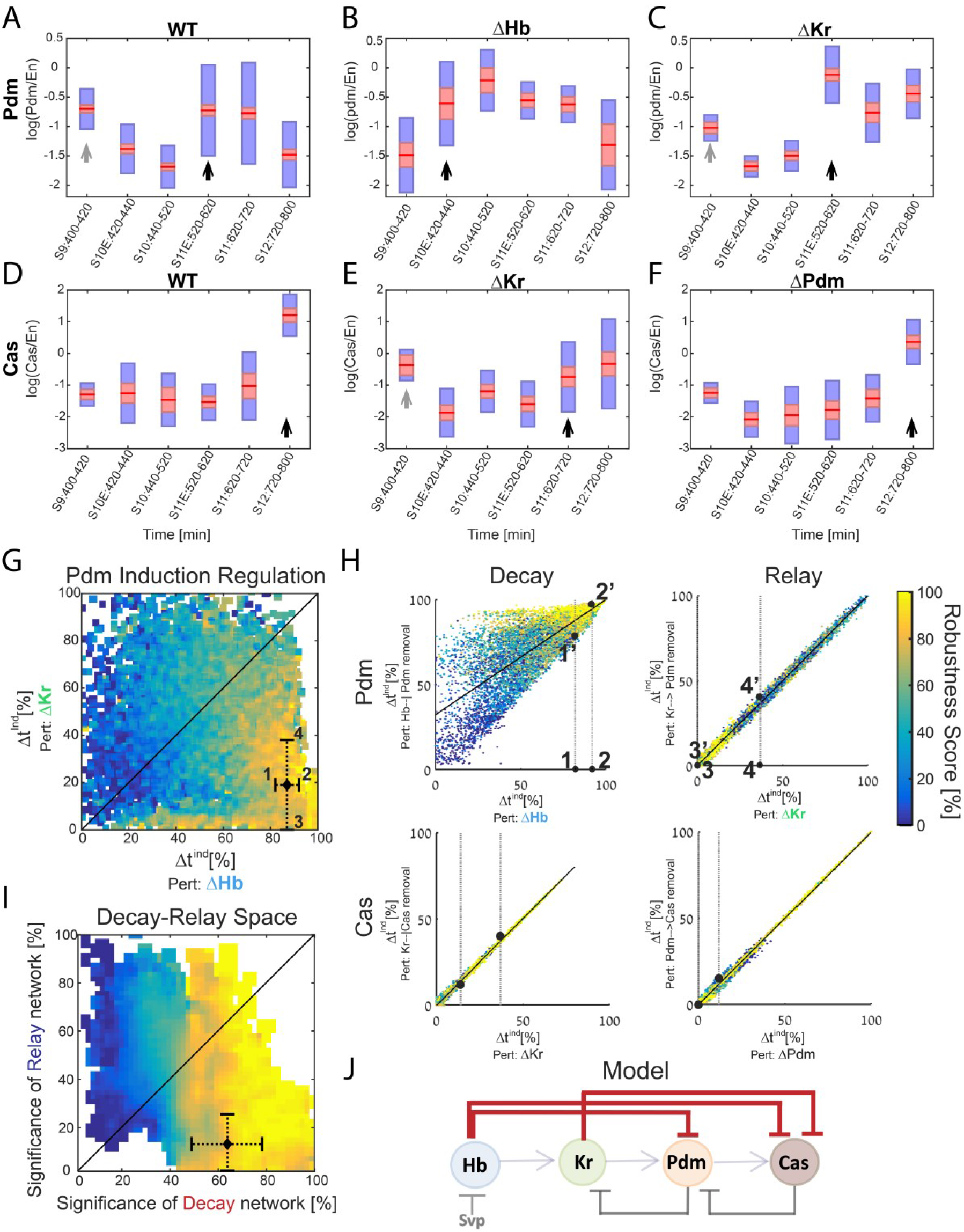
Positioning the in-vivo TTF circuit in the relay-520 decay timer space. (A-C) Pdm expression in NB7-1. Pdm expression was quantified in wild-type and mutant embryos. Data from the number of NB7-1 indicated in Table S2 is summarized by a box-plot, with 95% confidence interval shown in red and the 1 SD in blue. Developmental stages are indicated on the X axis. Stages of Pdm induction indicated by black arrows. Early transient inductions are indicated by grey arrows (see Methods for details). (D-F) Cas expression in NB7-1 wild type and mutant embryos. Data was plotted as described in (A-C). (G) Position of the *in-vivo* timer in the Pdm Hb-Kr sensitivity space: The measured changes in Pdm induction time following Kr and Hb deletions were used to position the *in-vivo* circuit in the Hb-Kr sensitivity space (black diamond), as described for the simulated data **Figure 3B**. Error bars are based on the experimental temporal resolution (see Methods for details). (H) Estimating the sensitivity of the *in-vivo* timer to removal of delay or relay interactions: The measured changes in Pdm and Cas induction times following TTF deletions were used to position the *in-vivo* circuit on the correlation plots from **Figure 3C-F** by taking the measured range denoted by 1,2 for ΔHb and 3,4 for ΔKr in **Figure 5G** and measuring the corresponding ranges for regulation removal. These are denoted by 1′,2′ and 3′, 4′ respectively in **Figure 5H** upper plots. Similarly, this was done for Cas regulation in **Figure 5H** lower plots (based on measured ranges in **Figure S1**, see Methods for details). (I) The TTF circuit is positioned in the region of repressor-delay timers: Data from A-H above defined the positioning of the TTF circuit on the decay-relay timer space (see text and Methods). Location of the in-vivo circuit is indicated by a black diamond, with error margins indicated by dashed error bars. (J) Our analysis indicates that the progress of TTF expression is dominated by repressor-decay, and not activator-accumulation. The NB TTF timer is shown again, with dominant decay regulations in bold.

Our measurements defined the change in Pdm and Cas induction times following the deletion of their activator or repressor TTFs. Both induction times showed higher sensitive to repressor deletion that to the deletion of their activator. These measurements allowed us to position the *in-vivo* circuit on the delay-relay space, **(c.f. Figure 1H)**. First, to estimate the robustness of the *in-vivo* circuit, we considered Pdm induction times, and positioned the *in-vivo* circuit on the Hb-Kr sensitivity space **(Figure 3B)** using the measured changes in Pdm induction time following deletion of Hb and of Kr. As can be seen, the *in-vivo* circuit was positioned in the narrow region in which robust circuits are found, strongly supporting the notion that the *in-vivo* circuit is indeed robust.

Second, we used our data of TTF deletion phenotypes to estimate the consequences of specifically removing the activator-relay or repressor-decay link. To this end, we used the tight correlations between the respective phenotypes observed in our simulations, as described in **Figure 3C-F**. Using these values to position the *in-vivo* circuit on the relay-decay space, clearly identified this circuit as a decay timer as was predicted by the robustness hypothesis. We therefore conclude that the timing of TTF expression is driven by a repressor-decay mechanism, rather than an activator-accumulation mechanism **(Figure 5G-I)**.

## Discussion

Our study suggests that a repressor-decay timer drives the sequential TTF expression to generate stereotyped temporal fate specification in *Drosophila* embryonic NB lineages **(Figure 5J)**. This finding may appear surprising, as previously this timer was thought to progress through a relay of activators achieved by feed-forward activation combined with feedback repression. A similar activator-relay mechanism was implicated also in driving the TTF cascade in *Drosophila* optic lobe NB lineages (Bertet et al., 2014; Li et al., 2013). Still, experimentally established crossregulation between the TTFs are consistent with both activator-relay and repressor-decay mechanisms, and the parameters defining the relative contributions of repressor decay or activator accumulation to the expression timing of each TTF were unknown.

We predicted that repressor-decay dominates TTF progression based on our computational results showing that this timer better buffers variation (‘noise’) in molecular parameters. Robust TTF progression is needed to maintain synchrony with the NB division cycles, a prerequisite for a reproducible NB lineage. At first sight, repressor degradation and activator accumulation may appear equivalent for measuring time delays. However, closer examination shows that they are in fact very different. First, activator accumulation requires continuous transcription while repressor degradation occurs following transcription shutdown. Second, activator accumulation approaches some steady state, which limits the possible readout thresholds. Furthermore, most of the dynamics is spent close to this threshold, so that small changes in threshold levels are translated into large changes in the measured delay time. By contrast, there is no such (theoretical) restriction on the readout threshold as repressor decays to zero expression. Together, these properties lead to different buffering capacities, both when considering a single-step timer, or in the context of the full TTF timer model. In all cases, encoding time-delays by repressor decay greatly promotes robustness.

Experimentally, we tested whether TTF progression is dominated by the repressor-decay interactions by measuring the expression timing of the last two TTFs in the cascade, Pdm and Cas. In both cases, TTF induction time was defined by the reduction in upstream repressor level, but showed little, if any change when upstream activator was deleted. Kr was not included in this analysis since we found that Kr was maintained in Hb mutant embryos (data not shown, (Isshiki et al., 2001)), indicating that Kr is induced by a factor external to the cascade, similarly to Hb. Notably, Kr remained constitutively expressed in embryos which were forced to express constitutive levels of Hb. Therefore, while Hb is an activator of Kr, it affects Kr expression not by determining its induction time but rather by determining shut-off time, allowing Kr decay only when Hb decays below a threshold, again implementing a repressor-decay, rather than an activator-relay timer. Hb is rapidly degraded during early embryogenesis, with an estimated half-life of ˜15 min (Okabe-Oho et al., 2009). While this half-life was not measured directly in NBs, it is likely to be similarly short based on the 1:1 relationship between *hb* transcriptional activity (detected with an intron probe) and Hb protein levels (detected with an antibody) (Grosskortenhaus et al., 2005). Assuming that Hb mRNA is similarly fast degrading, and that Hb is transcribed for only one cell-cycle, we estimate that Pdm starts expressing when Hb levels reduce to about 1-10% of their maximal value.

Our quantification of Pdm expression in wild-type embryos revealed that this phase is longer than previously thought, and led to the identification of a previously unrecognized Kr+Pdm+ GMC generated during this phase. Six motor neurons have previously been reported for the NB7-1 lineage (Landgraf et al., 1997; Schmid et al., 1999), yet there are only five Eve+ motor neurons in the lineage, raising the possibility that this “new” GMC may produce an Eve-negative motor neuron. Our analysis also revealed that Pdm is expressed in a burst during the Hb window. This was noted by Isshiki et al. (2001) but neither the functional significance nor the mechanism was discussed. Regarding function, we suggest that this early window of Pdm expression may allow it to be inherited in the first-born GMC, where in at least one lineage (NB4-2) it is required to specify first-born GMC identity, together with Hb (Bhat et al., 1995; Bhat and Schedl, 1994; McDonald et al., 2003; Yang et al., 1993; Yeo et al., 1995). Regarding mechanism, early Pdm and Cas expression is likely to be due to independent transcriptional activation of both genes, followed by repression of *pdm* transcription by Hb protein (Kambadur et al., 1998). The Pdm protein produced from the initial transcriptional burst, prior to Hb-mediated transcriptional repression, may persist into the new-born GMC. Alternatively, there may be a mechanism for blocking Hb repression of *pdm* transcription specifically in early-forming NBs.

In conclusion, we propose that the need to maintain robust gene expression timing within a noisy biological environment favored evolution of repressor-decay regulatory circuits controlling developmental patterning. This was previously shown for circuits that coordinate spatial patterning through the establishment of morphogen gradients or the control of direct cell-to-cell communication (Barkai and Shilo, 2009; Eldar et al., 2002; Eldar et al., 2003; Gavish et al., 2016; Rahimi et al., 2016). Our study suggests that robustness also played a major role in the design of developmental timers that function in neuronal differentiation.

## Materials and Methods

### Computational Methods

Randomized parameter sets (circuits) were generated by randomly selecting values for model parameters (See SI) from ranges indicated in **Table S3**. Parameters sere then substituted into model equations (See SI) and solved numerically by a standard MATLAB ODE solver. The solutions were tested for consistency: a consistent solution is one in which the temporal sequence of “on” (above threshold) TTFs is according to experimental observations for both WT and all mutants **(Figure 1C,D)**. Consistent parameter sets were scored for robustness to TTF production rates. When testing set robustness, we solved all combinatorial combinations of adding or subtracting 20% to all the TTFs production rates and then compared phase durations of each such noise combination set to those of the original set solution. Only if a noise combination yielded phase durations which are all within 10% distance of the respective original durations, the noise combination was considered “close” to the original. A robustness score was then calculated as the percentage of “close” noise combinations. The phases considered for this purpose were expression/co-expression phases leading to **different** neuronal fates **(Table S1)**. For example, a phase of Hb only expression followed by co-expression of Hb and Kr was considered a single phase since both lead to the 1 and 2 neuronal fates rendering the timing of Kr induction irrelevant in terms of NB lineage. In order to create perturbed parameter sets, parameter values or terms in model equations were changed accordingly: for TTF deletion the respective production rates were set to 0. For constitutive expression of a TTF, all the terms in model equations regulating this TTFs production were set to 1. For specific regulation removal, the regulation term was set to 1 (See SI for model equations). For **Figure 1H** and **Figure 2E**, significance scores of decay and relay for each consistent parameter set were first calculated with respect to Pdm and Cas separately. These scores were based on change in Pdm or Cas induction times caused by removing the decay or relay regulations governing these inductions. The difference in induction time for a perturbed set was calculated as
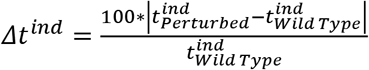 for each parameter set and then normalized as percentage out of the maximal *Δt^ind^* observed for all parameter sets, for this specific perturbation. The decay significance score, taking into account both Pdm and Cas induction regulation, was calculated by summing *Δt*^*ind*^ for Pdm (perturbation: removal of Hb--| Pdm) and *Δt*^*ind*^ for Cas (perturbation: removal of Kr--| Cas) and normalizing as percentage out of the maximal sum observed for all sets. The relay significance score was calculated similarly, only with relay removal perturbations. For **Figure 5G** and **Figure S1**, the calculation of *in-vivo* system location according to its *Δt*^*ind*^ for TTF deletion perturbations was performed by assuming the experimentally observed 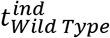 and 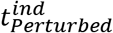 were the middle of the stage in which induction occurred, with an overall error margin of half that stage duration. These error margins were further increased when translating from *Δt*^*ind*^ for TTF deletions to *Δt*^*ind*^ for appropriate regulation removal. This translation was performed by placing the measured *Δt*^*ind*^ for TTF deletion on the correlation plots in **Figure 5H** and defining the range for regulation removal *Δt*^*ind*^ as the maximal range on the Y axis reached by robust (robustness score>80) sets within the X axis range of measured *Δt*^*ind*^ and its error margins. For **Figure 5I**, the joint Pdm and Cas decay and relay significance scores were again calculated by adding the scores for both TTFs and normalizing as previously described for simulated parameter sets. Normalization of measured *Δt*^*ind*^ for Pdm decay and relay removal was done by dividing by greatest experimentally observable values: Pdm induction at t=0 which corresponds to *Δt*^*ind*^ = 100% in the decay case and the middle of the last stage in the experiment (S12) in the relay case. For Cas, there was no need to normalize *Δt*^*ind*^ for relay removal since Cas induction in this case occurred during the last stage (S12) in the experiment. Cas decay removal *Δt*^*ind*^ was normalized by greatest *Δt*^*ind*^ observed in simulated parameter sets since we cannot expect Cas upregulation at t=0 because of inhibition by Hb.

### Experimental Methods

The following flies were used: (1) *y*^1^*W*^1^ (FBst0001495); (2) *hb*^*FB*^,*hb*^*p1*^/*TM3 ftz-lacZ* (Isshiki et al., 2001); (3) *Kr*^*CD*^,*Kr*^1^/*CyO wg-lacZ* (Isshiki et al., 2001); (4) *Df(2L)ED773/CyO wg-lacZ* (Grosskortenhaus et al., 2006); (5) *ac-VP16*^*AD*^,*gsb-Gal4*^*DBD*^ (Kohwi and Doe, 2013); (6) *w*^*1118*^; *10xUAS-IVS-myr::sfGFP-THS-10xUAS(FRT.stop)myr::smGdP-HA(attP2)(FBst0062127)*. Embryos were collected and incubated at 25°C until designated stages, and then fixed and stained with antibodies by following published protocols (Grosskortenhaus et al., 2005; Kohwi and Doe, 2013; Tran and Doe, 2008). The primary antibodies used in the studies were: rabbit anti-Ase (Cheng-Yu Lee, University of Michigan), mouse-anti-beta-galactosidase (Promega), rabbit anti-Cas (Mellerick et al., 1992) (Doe lab), rat anti-Dpn (Abcam, Eugene, OR), mouse anti-En 4D9 (Developmental Studies Hybridoma Bank (DHSB), Iowa City, IA), mouse anti-Hb (Abcam, Eugene, OR), guinea pig anti-Kr (Doe lab), rat anti-Pdm2 (Abcam, Eugene, OR), and rabbit anti-Wor (Doe lab). Fluorophore-conjugated secondary antibodies were from Jackson ImmunoResearch. Confocal images were taken by Zeiss LSM710 and protein quantities were measured with open software FIJI (Schindelin et al., 2012). Data was processed and plotted by Matlab. Mean volume and Standard deviation (STD) for all wild type NB7-1s were calculated. All NBs whose volume was further than 2 STD from mean volume (above or below) weren’t included in analysis, assuming these are currently dividing or miss identified cells. In order to determine the stage of Pdm and Cas induction from box plots in **Figure 5A-F**, a “background” level for both TTFs was defined as the mean of their mean levels (red line in box plots) in the second and third stages (S10E and S10), in WT. The first stage (S9) was not considered for this purpose due to possible early transient induction. The stage of induction was then defined as the first stage for which the 1.96 SEM (95% confidence interval-red rectangle) was above this mean, and was followed by an additional stage that satisfied this condition. The latter condition was disregarded for induction at the last stage of the experiment, stage S12.

## Acknowledgements

We thank Benny Shilo, Eyal Schejter, Danny Ben-Zvi and Gat Krieger for comments on the manuscript; Cheng-Yu Lee, Ward Odenwald and DHSB for antibodies. S.-L.L. and C.Q.D. were funded by the Howard Hughes Medical Institute and NIH R01-HD27056. N.B. was supported by a grant from the ERC.

## Author contributions

(A) I.A. did the modeling and conceived the project; S.-L.L. did the animal experiments; N.B. and C.Q.D. helped design experiments. All authors contributed to writing the manuscript.

## Supplemental Information

**Figure S1:**
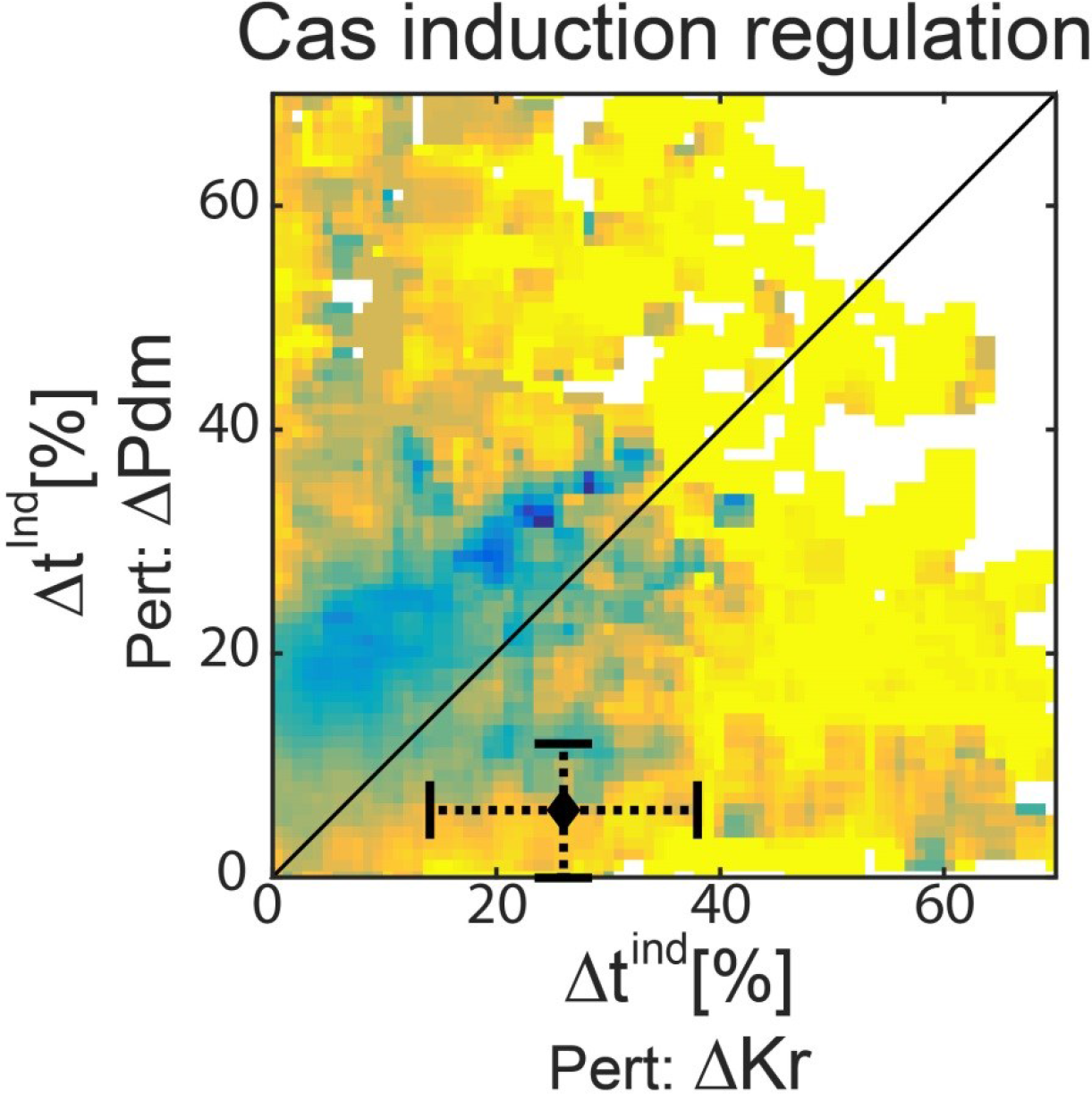
All consistent circuits, as described in Figures 1–2 above, were considered. Each consistent circuit was scored by measuring the change in Cas induction times following deletion of Kr or deletion of Pdm. These values were used to uniquely position each circuit in the Kr-Pdm sensitivity space. Color-coding circuits based on their robustness score, as in Figure 2E, shows that robust circuits are more often found in the region in the Kr-Pdm sensitivity space, in which Cas induction time is more sensitive to Kr deletion. Position of the in-vivo timer in the Cas Pdm-Kr sensitivity space: The measured changes in Cas induction time following Kr and Pdm deletions were used to position the in-vivo circuit in the Pdm-Kr sensitivity space, as described for the simulated data Figure 3B (Black diamond). Error bars are based on the experimental temporal resolution (see Methods for details).

**Table S1.** Neuronal fates induced by TTF co-expression in the NB. Neuronal fates for co-expression of TTFs in the NB at time of division were deduced from lineages described in Figure 1D. For every combination, the resulting fate is specified along with the genotypes from Figure 1D from which fate was deduced. Constitutive expression genotypes are denoted by const and deletions by Δ

| TTF combination | Neuronal fate | Reference genotype |
| --- | --- | --- |
| Hb | 1,2 | WT, const Hb, ΔHb |
| Hb,Kr | 1,2 | WT, const Hb, ΔHb |
| Kr | 3 | WT, ΔHb |
| Kr,Pdm | 3 | WT,const Kr |
| Pdm | 4 | WT, ΔPdm |
| Pdm,Cas | 5 | WT, ΔPdm, ΔCas,const Pdm |
| Cas | IN | const Cas |
| Hb,Pdm | 1,2 | const Pdm |

**Table S2.** Number of NB7-1s per box

| Figure, panel | Number of NB7-1s per box left to right |
| --- | --- |
| 4,C | 105,100 ,48, 81, 103, 90, 70,70,134,116, 63,77 |
| 4,E | 9,19,27,31,28,20,7,3 |
| 5,A | 105,100, 129 ,263 ,320 ,140 |
| 5,B | 31,28,22,26,22,16 |
| 5,C | 20,19,40 ,81 ,34 ,34 |
| 5,D | 19,35,18,38,28,35 |
| 5,E | 9,39,36,36,47,55 |
| 5,F | 17,27,27,40,27,44 |

## Model equations and Parameters

The molecular interactions driving the embryonic NB timer have been well described (Grosskortenhaus et al., 2005; Grosskortenhaus et al., 2006; Isshiki et al., 2001; Tran and Doe, 2008). These interactions appear to integrate the two core timers discussed in the main text: some genetic interactions are compatible with the decay-based timer, while other interactions are compatible with relay-based timer. Based on this existing knowledge, we formulated a mathematical model capturing the described interactions. This model allows for variable influence of both timer types of based on parameter choice. The following ODEs describe our TTF timer model:

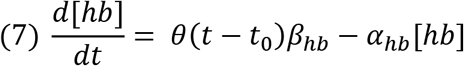

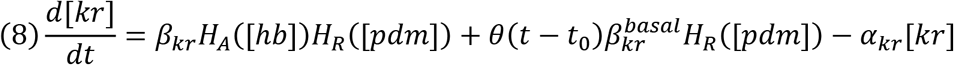

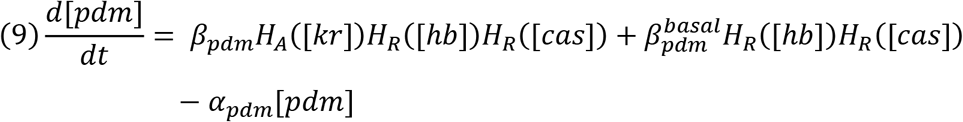

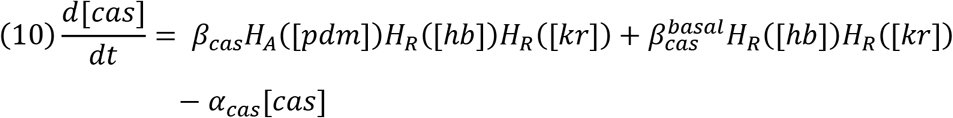

Notations and additional parameters:

*θ*(*t* - *t*_0_) it a temporal step function which allows the production of hb only at *t*<*t*_0_.
For the TTF i, *β*_*i*_ is i’s production rate, *α*_*i*_ is i’s degradation rate, 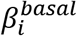 is i’s basal production rate in the absence of any activators.
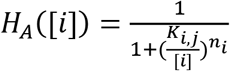, is a hill function representing transcriptional activation by the TTF i of gene j. With *K*_*i,j*_, the K_D_ for i activity on j, and *n*_*i*_, i’s hill coefficient.
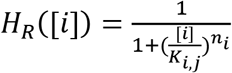, is a hill function representing transcriptional repression by the TTF i.

With ***K*_*i*_**, the K_D_ for i activity, and ***n*_*i*_**, i’s hill coefficient. Additional model parameters are the ***Tr*_*i*_**, which are the thresholds. Only when the concentration of i, [i], is above ***Tr*_*i*_**, i is considered to be “on”. **t_end_** is the duration of the simulation. We solve this full set of ODEs numerically using a standard MATLAB ODE solver. The model equations insure the dominance of the repressors: a gene will not be expressed in the presence of its repressor even if an activator is also present. This property which stems from the multiplication of the production rates by the ***H*_*A*_([*i*])*H*_*R*_([*i*])** term was observed experimentally (Grosskortenhaus et al., 2005; Grosskortenhaus et al., 2006; Isshiki et al., 2001; Nakajima et al., 2010; Tran and Doe, 2008; Tran et al., 2010).

**Table S3.** Parameter ranges used when searching for consistent sets. Drawing was done from a log-uniform distribution on indicated ranges. When no specific TTF is indicated for the parameter, it is the same for all four TTFs

| Parameter | Units | Range or values |
| --- | --- | --- |
| $t_0$ | $t[\text{min}]$ | 20 or 50 |
| $t_{end}$ | $t[\text{min}]$ | 280 |
| $\beta_i$ | $t^{-1}C$ | [1, 1000] |
| $\beta_{kr}^{basal}$ | $t^{-1}C$ | 0 for WT and all perturbations except Hb deletion. [1, 1000] for Hb deletion |
| $\beta_i^{basal}, i \in \{Pdm, Cas\}$ | $t^{-1}C$ | [1,800] |
| $\alpha_i$ | $t^{-1}$ | [0.1,10] |
| $K_{i,j}$ | $C^{n_w}$ | [0.1,2] |
| $Tr_i$ | C | 2 |
| $n_i$ | - | 5 |

